# A vitamin D-RelB/NF-κB pathway limits Chandipura virus multiplication by rewiring the homeostatic state of autoregulatory type 1 interferon-IRF7 signaling

**DOI:** 10.1101/2021.11.01.466649

**Authors:** Yashika Ratra, Naveen Kumar, Manti K. Saha, Chandrima Bharadwaj, Chen Chongtham, Sachendra S. Bais, Guruprasad Medigeshi, Gopalakrishnan A. Arimbasseri, Soumen Basak

## Abstract

Besides its functions in the skeletomuscular system, vitamin D also promotes protective immunity against viral pathogens. Viral sensing by mammalian cells triggers nuclear activation of RelA/NF-κB and IRF3 factors, which collaborate in mediating the early induction of antiviral type 1 interferons (T1-IFNs). Autocrine T1-IFN signaling further accumulates otherwise negligibly expressed IRF7 in virus-infected cells that then sustains T1-IFN production in a positive feedback. Surprisingly, prior cell-culture studies revealed that vitamin D actually suppresses signal-induced RelA activation. Indeed, it remains unclear how vitamin D limits viral multiplication in a cell-autonomous manner. Here, we examined the role of vitamin D in controlling cellular infections by the Chandipura virus (CHPV), a cytoplasmic RNA virus implicated in human epidemics. We found that vitamin D conditioning produced an altered cell state less permissive for CHPV multiplication because of the heightened expression of T1-IFNs. It is thought that viruses also induce a distinct RelB/NF-κB activity, which counteracts RelA-driven T1-IFN expressions in infected cells. Our analyses instead characterized a basal nuclear RelB activity, which was downregulated upon vitamin D-mediated suppression of RelB synthesis. Interestingly, this vitamin D-RelB pathway provoked IRF7-mediated positive autoregulation augmenting constitutive T1-IFN expressions even in the absence of viral infections. Accordingly, RelB deficiency rendered redundant, while IRF7 depletion abrogated antiviral vitamin D actions. In sum, our study suggests that the homeostatic state of the signaling circuitry comprising of the NF-κB and T1-IFN pathways connects micronutrients to antiviral immunity at the cellular level.

**Significance statement:** Vitamin D limits viral infections, but the underlying mechanism remains unclear. Linking micronutrients to antiviral immunity, Ratra et al. characterize an immune signaling circuitry engaged by vitamin D that generates a cellular state less permissive to infections by Chandipura virus, a pathogen of public health importance.

## Introduction

Besides its well-documented role in the skeletomuscular system, vitamin D also modulates protective immunity against viruses (Siddiqui et al., 2020). Population-level observational studies argued that vitamin D deficiency elevates the risk of lung infection by the respiratory syncytial virus (RSV) and influenza A virus and exacerbates inflammatory lung injuries (Belderbos et al., 2011; Greiller and Martineau, 2015; Sabetta et al., 2010). Similarly, a weakened vitamin D pathway escalated hepatitis C virus infections and the incidence of hepatocellular carcinoma (Jin et al., 2018). Vitamin D deficiency prolonged inflammatory polyarthritis in Chikungunya virus (CHIKV)-infected individuals (Bhavana et al., 2016). On the other hand, dietary vitamin D supplementation significantly diminished influenza A virus replication and downregulated pro-inflammatory cytokines in the infected mice (Hayashi et al., 2020). Ex vivo vitamin D treatment of airway epithelial cells restricted inflammatory gene expressions upon RSV infection (Hansdottir et al., 2010). Vitamin D also diminished rotavirus infection in vivo and ex vivo (Zhao et al., 2019). Although randomized control trials supplementing vitamin D are yet to produce encouraging outcomes in human subjects, monocyte-derived macrophages from vitamin D supplemented individuals were refractory to dengue virus infection and produced a reduced level of inflammatory mediators (Giraldo et al., 2018). These studies establish that vitamin D restricts the multiplication of a broad spectrum of viral pathogens and limits infection-inflicted inflammatory pathologies.

Viral invasion of tissue-resident cells triggers the nuclear accumulation of the RelA transcription factor *via* the canonical NF-κB pathway (Bowie and Unterholzner, 2008). RelA promotes the expression of pro-inflammatory cytokines and chemokines, which recruit macrophages to the infected site, and instruct effector T cells. While cytotoxic T lymphocytes are critical for eliminating virus-infected cells, Th1 and Th17 CD4 T cell subsets further fuel local inflammation, often causing tissue injuries. In collaboration with another virus-activated factor IRF3, RelA also mediates the early induction of T1-IFNs. T1-IFNs engage in autocrine and paracrine signaling through the cognate T1-IFN receptor (IFNAR) that limits viral replication in infected cells and impedes infection of neighboring cells by progeny virus particles. Notably, T1-IFN signaling promotes cellular accumulation of otherwise negligibly expressed IRF7, which sustains T1-IFN production in positive feedback (Ning et al., 2011; Sato et al., 1998). In addition, a separate noncanonical NF-κB pathway promotes nuclear translocation of RelB, which directs naïve lymphocytes to secondary lymphoid organs by upregulating homeostatic chemokines in stromal cells (Mukherjee et al., 2017; Sun, 2017). Consistent with RelB’s role in the adaptive immune compartment, RelB deficiency in mice severely compromised cytotoxic T cell responses against lymphocytic choriomeningitis virus (Weih et al., 1997). Interestingly, it has also been reported that virus infections stimulate the noncanonical RelB/NF-κB pathway and that virus-induced noncanonical RelB activity inhibits the expression of antiviral T1-IFNs (Jin et al., 2014). Thus, an intricate cell signaling circuitry appears to calibrate immune responses against viral pathogens.

Vitamin D tunes these immune regulatory pathways. For instance, engagement of vitamin D receptor (VDR) impedes signal-responsive activation of the inflammatory, canonical RelA/NF-κB pathway (Chen et al., 2013). In T cells, vitamin D downregulates the transcription of the genes encoding the Th1 cytokine IFNγ (Cippitelli and Santoni, 1998) and the Th17 cytokine IL-17 (Joshi et al., 2011). More so, vitamin D supports suppressive regulatory T cells in various anatomic niches (Joshi et al., 2011). These studies provide an explanation of how vitamin D mitigates pathological inflammation induced by microbial entities, including viruses (Wei and Christakos, 2015). Vitamin D treatment of cells also upregulates the expression of the antimicrobial peptide cathelicidin (Wang et al., 2004). However, cathelicidin inhibits mainly bacterial pathogens, and its antiviral functions are rather selective (Wei and Christakos, 2015). Indeed, it remains enigmatic how vitamin D exerts a broad cell-autonomous antiviral function while also negatively impacting T1-IFN-inducing RelA activity.

Chandipura virus (CHPV) is a neurotropic RNA virus belonging to the Rhabdoviridae family and vesiculovirus genera (Basak et al., 2007). This virus has been implicated in several recent epidemic outbreaks of acute encephalitis in the Indian subcontinent that were characterized by influenza-like illness, neurological manifestations, and a high case fatality rate (Menghani et al., 2012). It has been suggested that inadequate peripheral immune reactions, including those elicited by cells of the monocyte and macrophage lineage, allow the dissemination of CHPV to the central nervous system triggering neuronal infection and widespread cell death (Verma et al., 2018). CHPV infection of cultured cells readily induces canonical RelA/NF-κB and IRF3 activities in the nucleus (Bais et al., 2019; Pandey et al., 2021). Therefore, a role of RelA-mediated gene expressions in aggravating CHPV-inflicted neuroinflammation cannot be ruled out. Nevertheless, recurrent outbreaks prompted global concerns leading to the recognition of CHPV as an important human pathogen. Despite widespread micronutrient deficiency in India and other developing countries (Ministry of Health and Family Welfare India), a plausible role of vitamin D in CHPV pathogenesis has not been studied.

Here, we investigated antiviral vitamin D functions in the context of cellular infections by two RNA viruses. We found that vitamin D conditioning triggered heightened expressions of T1-IFNs, leading to an altered cell state less permissive for CHPV and also CHIKV propagation. Our mechanistic studies revealed that vitamin D downregulated the expression, and consequently the basal activity, of RelB provoking the autoregulatory T1-IFN-IRF7 axis even in the absence of viral infections. Accordingly, IRF7 depletion abrogated, while RelB deficiency rendered redundant, the antiviral vitamin D actions. In sum, our study unveils a complex signaling circuitry comprising of the NF-κB and T1-IFN pathways that connects micronutrients to antiviral cellular homeostasis.

## Results

### Vitamin D conditioning produces an altered cell state less permissive for CHPV propagation

We set up ex vivo CHPV infection experiments for probing antiviral vitamin D functions. To this end, cultured human or mouse cells were treated for 24h with 100nM of 1α, 25-dihydroxyvitamin D3, the active form of vitamin D, and then these vitamin D-conditioned cells were infected either in the continuing presence of vitamin D or subsequent to withdrawal of vitamin D from the culture medium. As a control, vitamin D naïve cells were used. At 12h post-infection, cells were harvested for biochemical analyses, and the culture supernatant was subjected to plaque assay for determining the abundance of progeny virus particles (Figure 1A, and also see Materials and Methods).

**Figure 1:**
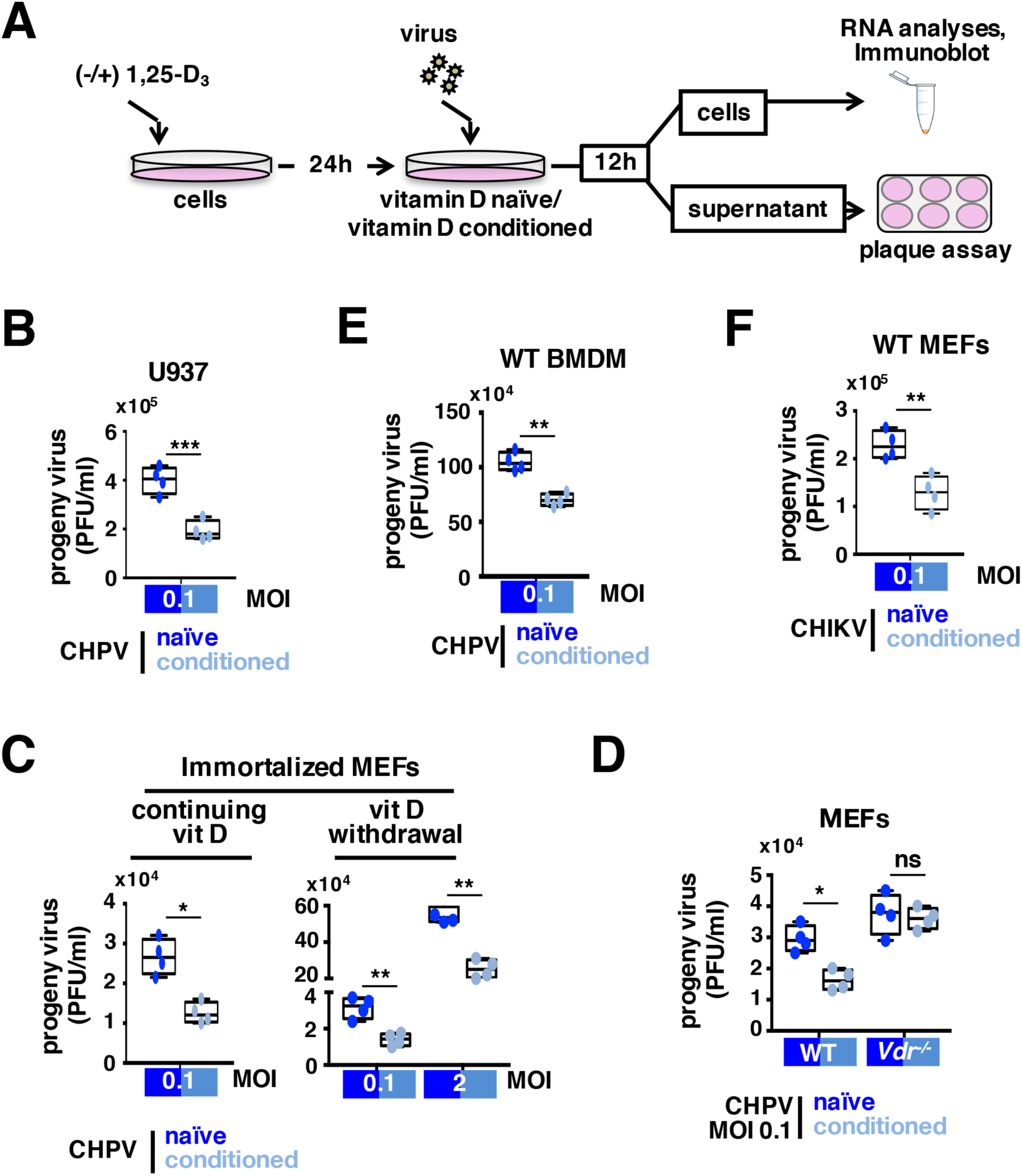
Cell conditioning with 1, 25-dihydroxyvitamin D3 diminishes the progeny yield upon CHPV infection. **A**. A schematic presentation of the cell treatment and infection regime. Briefly, cells were treated with 1α, 25-dihydroxyvitamin D3 for 24h, and then vitamin D-conditioned cells were infected with virus subsequent to vitamin D withdrawal. For certain specified sets, conditioned cells were infected and cultured in the continuing presence of vitamin D. As a control, vitamin D naïve cells were used. The culture supernatant was collected at 12h post-infection and subjected to plaque assay. Infected cells were also harvested at various times post-infection for protein and RNA analyses. When used, immortalized MEFs were specifically identified in the figure labels. **B**. and **C**. Plaque assay revealing the titer of progeny virus produced upon infection of vitamin D-conditioned U937 cells **(B)** or immortalized WT MEFs **(C)** with CHPV at the indicated MOI either in the continuing presence of vitamin D **(Figure 1C, left panel)** or subsequent to vitamin D withdrawal **(Figure 1B; Figure 1C, right panel)**. **D**. and **E**. Vitamin D naïve or vitamin D-conditioned WT and *Vdr*^*-/-*^ primary MEFs were infected with 0.1 MOI of CHPV, and the progeny virus yield was determined by plaque assay **(D)**. Similarly, the effect of vitamin D conditioning of BMDMs on CHPV propagation was scored **(E)**. **F**. Barplot comparing the titer of progeny virus produced upon CHIKV infection of vitamin D naïve or vitamin D-conditioned WT primary MEFs. The data represent means ± SEM of four biological replicates. *, *P* ≤ 0.05; **, *P* ≤ 0.01; ns, not significant.

We found that vitamin D conditioning of human-derived monocytic U937 cells indeed caused a significant reduction in the progeny CHPV titer (Figure 1B). Because mouse-derived cell systems facilitate genetic dissection of immune signaling mechanisms, we then focused on mouse embryonic fibroblasts (MEFs), including those immortalized using the NIH 3T3 protocol. Regardless of the continuing presence of vitamin D, conditioning of immortalized MEFs led to more than two-fold decrease in progeny titer upon 12h of CHPV infection at MOI 0.1 (Figure 1C). Vitamin D withdrawal did not impact the ability of these conditioned cells to contain CHPV growth even when they were challenged at MOI 2. Therefore, in all our subsequent experiments, we infected vitamin D-conditioned cells with viruses after vitamin D withdrawal. Importantly, a comparison of primary MEFs derived from WT and *Vdr*^*-/-*^ mice ascertained that the observed vitamin D conditioning effect required functional VDR signaling (Figure 1D). As compared to MEFs, bone marrow-derived macrophages (BMDMs) generated from WT mice were more proficient in producing progeny particles upon CHPV infection (Figure 1E and Figure S1A). Vitamin D treatment substantially diminished CHPV propagation also in BMDMs. Furthermore, vitamin D conditioning of MEFs hindered the multiplication of CHIKV, another RNA virus of public health importance (Figure 1F). We infer that vitamin D produces a moderate but functionally significant inhibitory effect on CHPV as well as CHIKV propagation in diverse cell types of both human and mouse origin. Vitamin D conditioning also seems to impart a lasting antiviral cell state, reminiscing of innate immune memory, obviating the requirement of continuing vitamin D treatment.

### Vitamin D treatment potentiates the expression of antiviral T1-IFN genes

Particularly in ex vivo settings, T1-IFNs play a dominant role in controlling viral propagation. We asked if the engagement of T1-IFN signaling provides for the antiviral effect of vitamin D. To address this question, we subjected a previously published transcriptomic dataset generated using vitamin D naïve and vitamin D-conditioned THP1 monocytic cells to gene set enrichment analyses (GSEA) (Salamon et al., 2014). Our genome-scale investigation identified significant enrichment of T1-IFN-related genes among those upregulated in response to vitamin D treatment (Figure 2A). Three of the top ten most enriched GO for biological processes terms among genes induced in THP1 cells by vitamin D were linked to the antiviral T1-IFN pathway (Figure S1B). Furthermore, our RT-qPCR analyses involving WT MEFs disclosed that vitamin D conditioning led to an approximately two-fold increase in the basal expression of genes encoding IFNβ and IFNα4 (Figure 2B). CHPV infection more profoundly elevated the abundance of these mRNAs in vitamin D-conditioned MEFs as compared to naïve cells. Consistent with its role in upregulating T1-IFNs, vitamin D conditioning augmented the expression of interferon-stimulated genes, including those encoding ISG56, ISG15, and OAS1, in both uninfected and CHPV-infected MEFs (Figure 2C). As a control, we measured the level of IL-1β mRNA, whose expressions are known to be T1-IFN independent. We did not notice any discernible change in the abundance of IL-1β mRNA upon vitamin D treatment in either uninfected or infected cells. Importantly, strengthening T1-IFN signaling in vitamin D-conditioned MEFs correlated with a significantly reduced accumulation of CHPV genome RNA upon cell infection (Figure 2D). We identified that genetic deficiency of IFNAR expression not only amplified the production of progeny CHPV particles in infected *Ifnar1*^*-/-*^ MEFs but also blunted the antiviral effect of vitamin D conditioning (Figure 2D). We conclude that vitamin D conditioning directs T1-IFN signaling to impede CHPV propagation.

**Figure 2:**
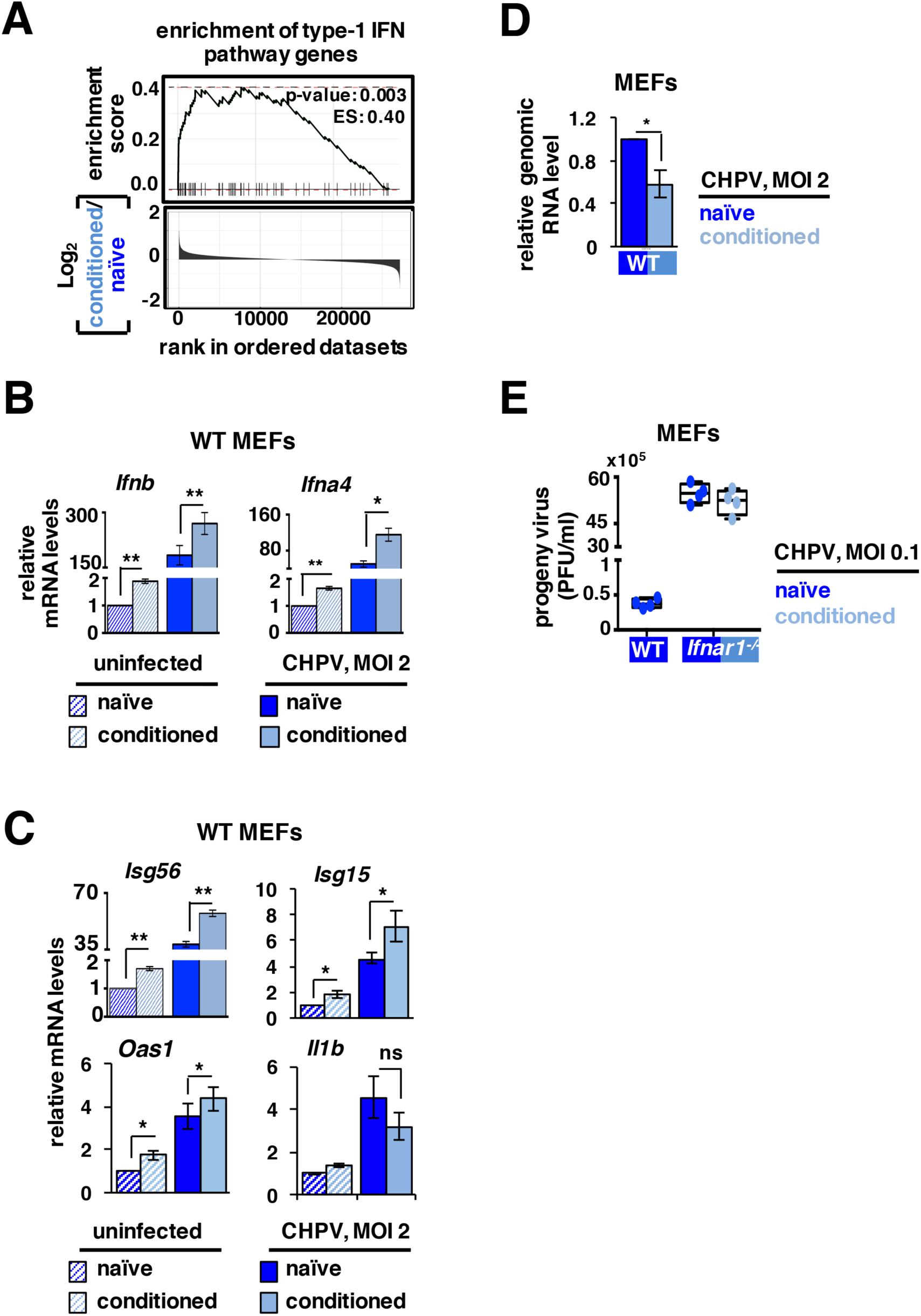
Vitamin D conditioning potentiates antiviral T1-IFN gene expressions. **A**. GSEA examining the impact of vitamin D on T1-IFN signaling. We utilized a dataset (accession no. GSE57028) describing vitamin D induced changes in the gene expression in THP1 monocytic cells. Subsequent to correcting for the detection p-value and signal to noise ratio, we ranked the genes in descending order based on the fold change difference of their expressions between vitamin D-conditioned and vitamin D naïve cells (bottom panel). The relative enrichment of T1-IFN pathway genes in this ranked list was determined by GSEA (top panel); each of the horizontal dashed line represents a T1-IFN pathway gene. **B**. and **C**. RT-qPCR revealing the relative abundance mRNAs encoding the indicated T1-IFNs **(B)** and ISGs **(C)** in vitamin D naïve or vitamin D-conditioned WT primary MEFs either left uninfected or infected with 2 MOI of CHPV for 12h. As a control, the abundance of IL-1β mRNA was also determined **(C)**. **D**. RT-qPCR showing the relative abundance of viral genome RNA in vitamin D-conditioned WT primary MEFs infected with 2 MOI of CHPV for 12h. **E**. Plaque assay scoring the titer of progeny virus produced by vitamin D naïve or vitamin D-conditioned *Ifnar1*^*-/-*^ primary MEFs upon infection with 0.1 MOI of CHPV. Vitamin D naïve, virus-infected WT MEFs were used as controls. The RT-qPCR data represent means ± SEM of three independent experiments. The plaque assay data represent means ± SEM of four biological replicates. *, *P* ≤ 0.05; **, *P* ≤ 0.01; ns, not significant.

### Vitamin D conditioning depletes nuclear RelB by downregulating RelB mRNA

RelA and IRF3 positively regulate the expression T1-IFNs, while RelB suppresses IFNβ production (see Introduction). To understand how vitamin D induced T1-IFNs, we probed the nuclear abundance of these factors. Our immunoblot analyses revealed that irrespective of vitamin D treatment, CHPV infection accumulated phosphorylated IRF3 protein in the nuclear extracts from WT MEFs (Figure 3A). Phosphorylated IRF3 was not detectable in the uninfected cells. Moreover, naïve cells possessed a low basal RelA activity in the nucleus that was intensified upon CHPV infection (Figure 3A), owing to virus-induced canonical signaling (Bais et al., 2019). We observed that vitamin D conditioning subtly impacted the RelA activity, both in uninfected and CHPV-infected MEFs. We further noticed a RelB activity in uninfected naïve cells that was modestly upregulated upon CHPV infection. Interestingly, vitamin D treatment drastically curtailed the basal RelB activity in the nucleus and also prevented its virus-induced nuclear accumulation (Figure 3A).

**Figure 3:**
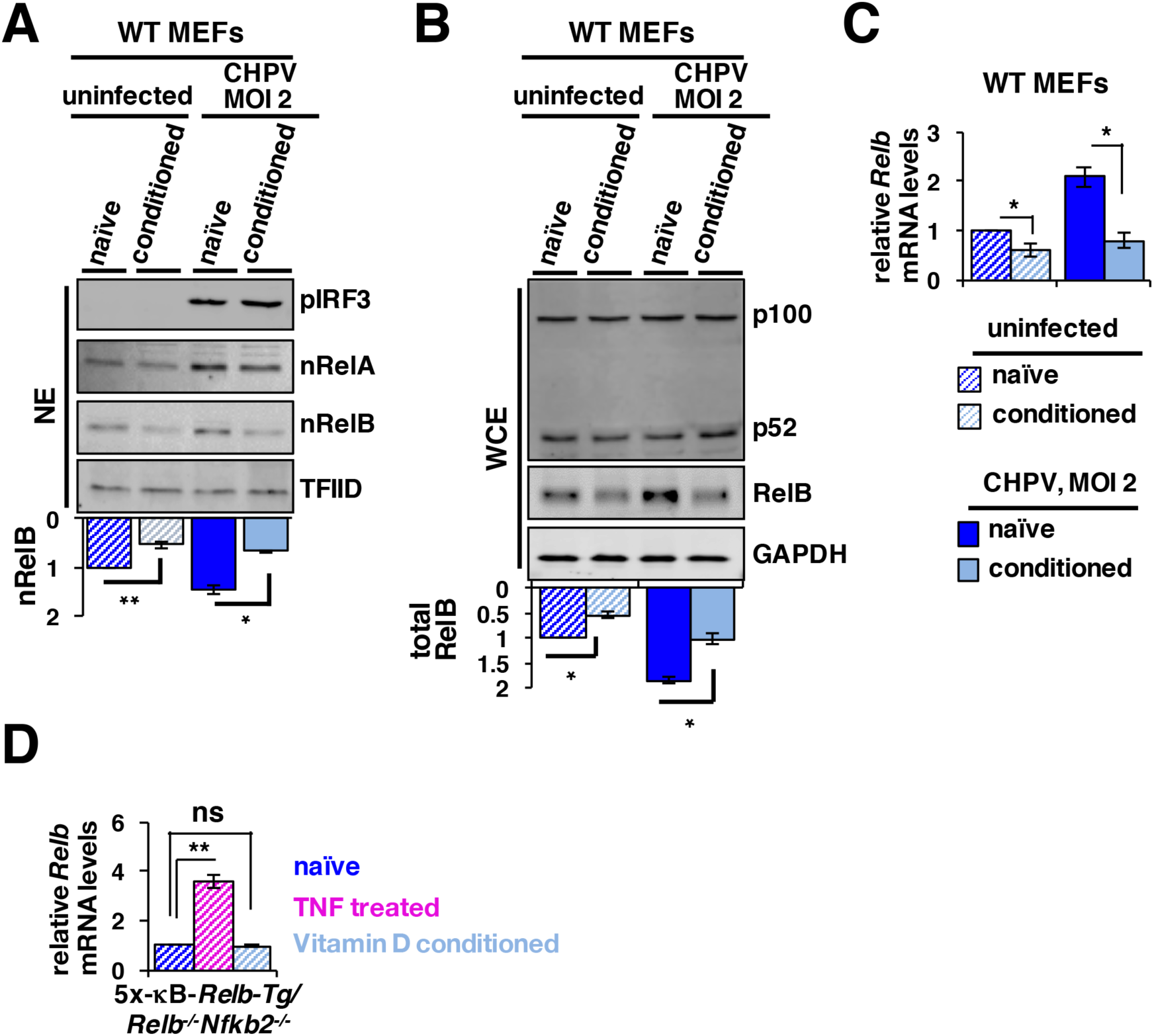
Cell conditioning with vitamin D reduces the nuclear abundance of RelB by repressing RelB mRNA expressions. **A**. and **B**. Immunoblot analyses of nuclear **(A)** and whole-cell **(B)** extracts derived from vitamin D naïve or vitamin D-conditioned WT MEFs either left uninfected or infected with 2 MOI of CHPV for 12h. TFIID and GAPDH served as loading controls for nuclear and cytoplasmic extracts, respectively. The signal corresponding to RelB was quantified from three independent experiments and presented below the respective panels. **C**. and **D**. RT–qPCR analysis revealing the relative abundance of RelB mRNA in vitamin D naïve or vitamin D-conditioned WT MEFs either left uninfected or infected with 2 MOI of CHPV for 12h **(C)**. The effect of vitamin D conditioning on the level of RelB mRNA was similarly captured using immortalized *Relb*^*-/-*^*Nfkb2*^*-/-*^ MEFs expressing RelB from a transgene (Tg) under a promoter consisting of five tandem κB sites **(D)**. TNF treated cells were used as controls. The quantified data represent means ± SEM of three independent biological experiments. *, *P* ≤ 0.05; **, *P* ≤ 0.01; ns, not significant.

In resting cells, RelB is mostly sequestered in the cytoplasm by the NF-κB precursor protein p100 (Sun, 2017). Noncanonical NF-κB signaling induces proteasomal processing of p100 into the mature NF-κB subunit p52, releasing the RelB:p52 dimer in the nucleus. We asked if noncanonical signaling promoted nuclear accumulation of RelB upon CHPV infection and was, in fact, targeted by vitamin D. Accordingly, we determined the activity of NIK, the noncanonical pathway-activating kinase, and scored the cellular abundance of p52 relative to p100. CHPV infection did not activate NIK or stimulate p100 processing to p52 in MEFs in our experiments (Figure 3B and Figure S1C). We instead noted a modest level of constitutive p100 processing in naïve cells that was also insensitive to vitamin D treatment. Curiously, CHPV infection of naïve cells increased the abundance of RelB protein in the whole-cell extracts that was accompanied by a close to two fold increase in the level of RelB mRNA (Figure 3B and Figure 3C). Vitamin D conditioning significantly diminished the level of cellular RelB protein as well as mRNA in both uninfected and infected cells. RelA is known to drive the expression from the *Relb* promoter, which harbors κB sites (Bren et al., 2001). We further examined the effect of vitamin D conditioning using *Relb*^*-/-*^*Nfkb2*^*-/-*^ MEFs expressing RelB from a retroviral transgene under a minimal NF-κB-responsive promoter consisting of five κB sites. We found that TNF treatment induced RelB mRNA production from this transgenic construct (Figure 3D). However, this minimal κB promoter was not sufficient for downregulating RelB expression in response to vitamin D. We deduce that CHPV infection promotes nuclear accumulation of RelB independent of the noncanonical NF-κB pathway, presumably involving enhanced RelB production and that vitamin D depletes nuclear RelB in both uninfected and infected MEFs by downmodulating RelB transcription involving RelA/NF-κB independent mechanisms.

### RelB restrains constitutive activation of autoregulatory type 1 IFN-IRF7 signaling

Next, we questioned if depletion of RelB was linked to T1-IFN upregulation. To this end, we compared WT and *Relb*^*-/-*^ MEFs in global gene expression studies. Our GSEA showed that T1-IFN pathway genes were significantly enriched among those basally upregulated in the absence of RelB (Figure 4A). Our RT-qPCR analyses confirmed a more than five-fold increase in the constitutive level of mRNAs encoding IFNβ and IFNα4 in *Relb*^*-/-*^ MEFs compared to WT cells (Figure 4B). Consistently, RelB deficiency led to vastly augmented basal expressions of ISG56, ISG15, and OAS1 mRNAs (Figure 4C). Except for ISG56, these mRNAs were also more abundant in CHPV-infected *Relb*^*-/-*^ cells (Figure 4B and Figure 4C). However, the difference in their levels between WT and *Relb*^*-/-*^ MEFs became less pronounced upon CHPV infection. Importantly, we observed a close to twenty-fold increase in the basal expression of the interferon-stimulated gene encoding IRF7 in RelB-deficient cells; the level of IRF7 mRNA was considerably higher also in CHPV-infected *Relb*^*-/-*^ MEFs (Figure 4C). As a control, we scored the abundance of RelA mRNA, which was equivalently expressed in WT and *Relb*^*-/-*^ cells. Our analyses ascertained that RelB particularly restrained the basal expression of T1-IFNs.

**Figure 4:**
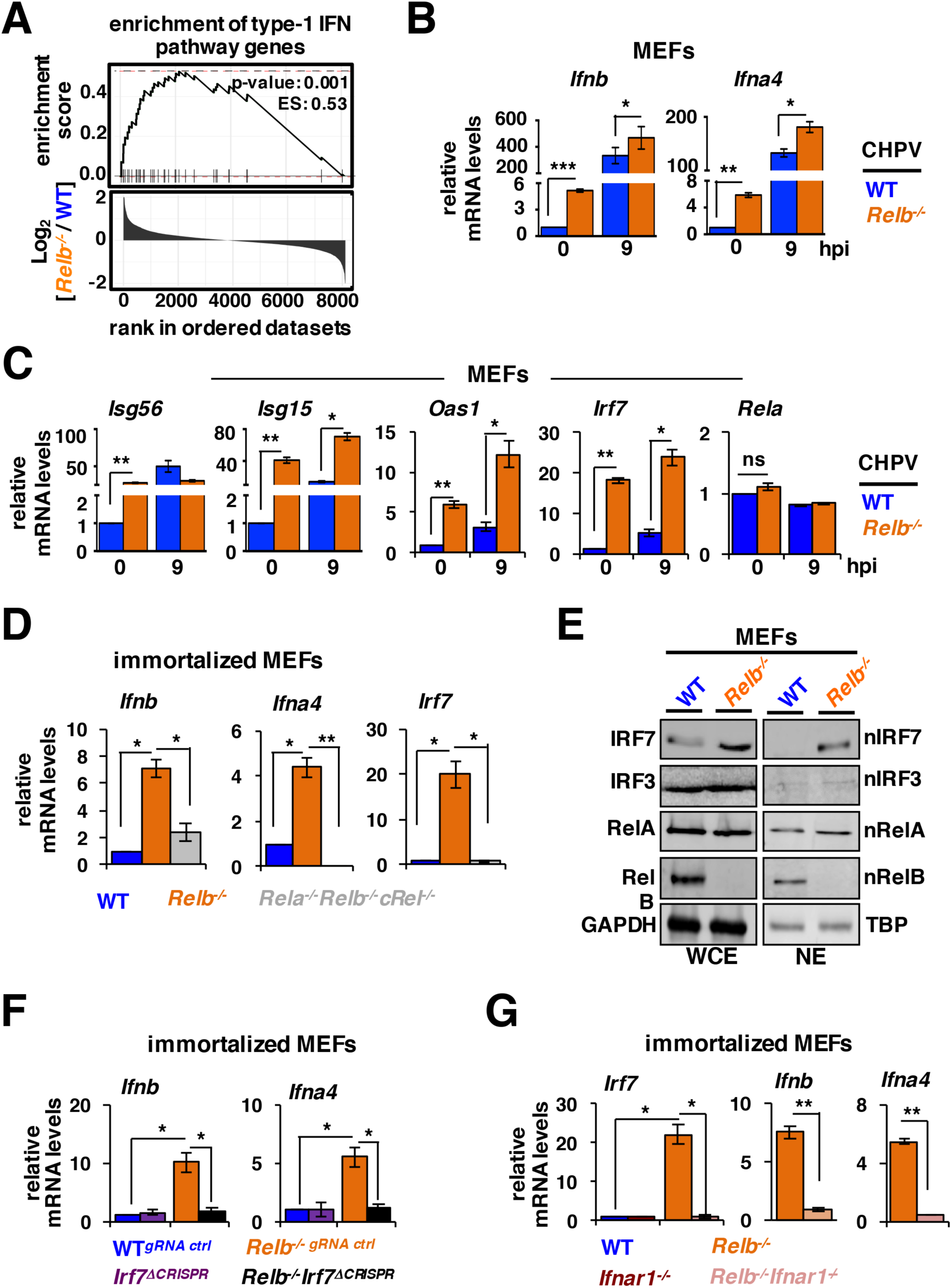
RelB deficiency triggers elevated basal expressions of T1-IFNs involving IRF7-mediated positive autoregulation. **A**. GSEA comparing WT and *Relb*^*-/-*^ MEFs for the basal expression of T1-IFN pathway genes. Genes were ranked based on the fold change difference in their basal expressions between *Relb*^*-/-*^ and WT MEFs. **B**. and **C**. RT-qPCR analyses comparing WT and *Relb*^*-/-*^ primary MEFs for basal and CHPV-induced expressions of genes encoding indicated T1-IFNs **(B)** and ISGs **(C)**. Cells were infected with 2 MOI of CHPV for 9h. **D**. and **G**. Immortalized MEFs of the indicated genotypes were subjected to RT-qPCR analyses for scoring basal expressions of IFNβ and IRF7 mRNAs. The results are relative to immortalized WT MEFs. **E**. Whole-cell (left panel) and nuclear (right) extracts from uninfected WT and *Relb*^*-/-*^ MEFs were subjected to immunoblot analyses using antibodies against the indicated proteins. **F**. RT-qPCR revealing basal expression of IFNβ and IFNα4 mRNAs in immortalized MEFs of the indicated genotypes. By subjecting WT and *Relb*^*-/-*^ immortalized MEFs to CRISPR-Cas9-mediated genome editing, we generated control *Irf7*^Δ*CRISPR*^ cells and *Relb*^*-/-*^*Irf7* ^Δ*CRISPR*^ cells, which lacked IRF7 expressions. The quantified data represent means ± SEM of three biological replicates. *, *P* ≤ 0.05; **, *P* ≤ 0.01; **, *P* ≤ 0.001; ns, not significant.

It was suggested that noncanonical RelB NF-κB signaling counteracts RelA-driven expression of T1-IFNs in virus-infected cells (Jin et al., 2014). We found that genetic deficiency of RelB instigated constitutively elevated T1-IFN signaling and IRF7 expression even in uninfected MEFs. We asked if RelA modulated the homeostatic state of the T1-IFN pathway in *Relb*^*-/-*^ cells. Our analyses using immortalized MEFs revealed a similar increase in the basal abundance of IFNβ, IFNα4 as well as IRF7 mRNAs in the absence of RelB (Figure 4D). Indeed, IFNβ and IFNα4 mRNA levels were restored to that of WT MEFs in *Relb*^*-/-*^ *Rela*^*-/-*^*cRel*^*-/-*^ cells, which additionally lacked RelA and cRel, constituents of the NF-κB dimers activated by the canonical pathway. Interestingly, this fall in the basal expression of T1-IFN cytokines in *Relb*^*-/-*^*Rela*^*-/-*^*cRel*^*-/-*^ cells was associated with a concomitant reduction in the IRF7 mRNA level (Figure 4D).

Because IRF7 and T1-IFNs are intimately linked through an autoregulatory loop, we examined if, in addition to RelA, IRF7 reinforced T1-IFN signaling in RelB-deficient cells. Corroborating gene expression studies (Figure 4C), our immunoblot analyses showed substantially increased accumulation of IRF7 in the whole-cell extracts from *Relb*^*-/-*^ primary MEFs (Figure 4E). Importantly, IRF7 could be easily detected in the nuclear extracts derived from *Relb*^*-/-*^, but not WT, cells in its basal state. WT and RelB-null MEFs were both devoid of the constitutive IRF3 activity in the nucleus, whereas RelB deficiency led to only a subtle enhancement of the constitutive nuclear abundance of RelA (Figure 4E). Remarkably, CRISPR-Cas9-mediated disruption of IRF7 expression completely abolished heightened basal synthesis of IFNβ and IFNα4 mRNAs in *Relb*^*-/-*^ cells (Figure 4F and Figure S1D). IRF7 deletion in WT cells had virtually no impact on constitutive T1-IFN expressions. We further generated immortalized *Relb*^*-/-*^*Ifnar1*^*-/-*^ MEFs for our experiments. Our mRNA analyses suggested that signaling through IFNAR was important for the elevated basal expression of IRF7 as well as T1-IFN mRNAs in the RelB-null genetic background (Figure 4G). We interpret that in the absence of RelB, RelA (and cRel) primes constitutive expression of IFNβ, which accumulates cellular IRF7 involving autocrine IFNAR signaling, and that IRF7 sustains heightened basal expression of T1-IFNs, including IFNα and IFNβ mRNAs, in RelB deficient cells involving positive autoregulatory loop.

### Autoregulatory type 1 IFN-IRF7 signaling limits viral growth in RelB-deficient cells

We then investigated if RelB-deficient cells restricted viral infections engaging autoregulatory T1-IFN-IRF7 signaling. Compared to WT cells, *Relb*^*-/-*^ MEFs produced approximately four-fold fewer progeny virus particles upon infection with CHPV at MOI 0.1; the difference was even more prominent when cells were infected at MOI 2 (Figure 5A). Our mRNA analyses consistently revealed a substantially less abundance of viral genomic RNA and mRNAs in *Relb*^*-/-*^ MEFs at 9h post-CHPV infection (Figure 5B). Likewise, RelB deficiency hindered CHPV propagation in BMDMs and immortalized MEFs (Figure 5C). In particular, immortalized *Relb*^*-/-*^ MEFs displayed a dramatic effect with close to eight- and eleven-fold reduction in the progeny particles upon CHPV infection at MOI 0.1 and MOI 2, respectively (Figure 5C, right panel). Moreover, *Relb*^*-/-*^ MEFs were also less supportive for CHIKV growth (Figure 5D).

**Figure 5:**
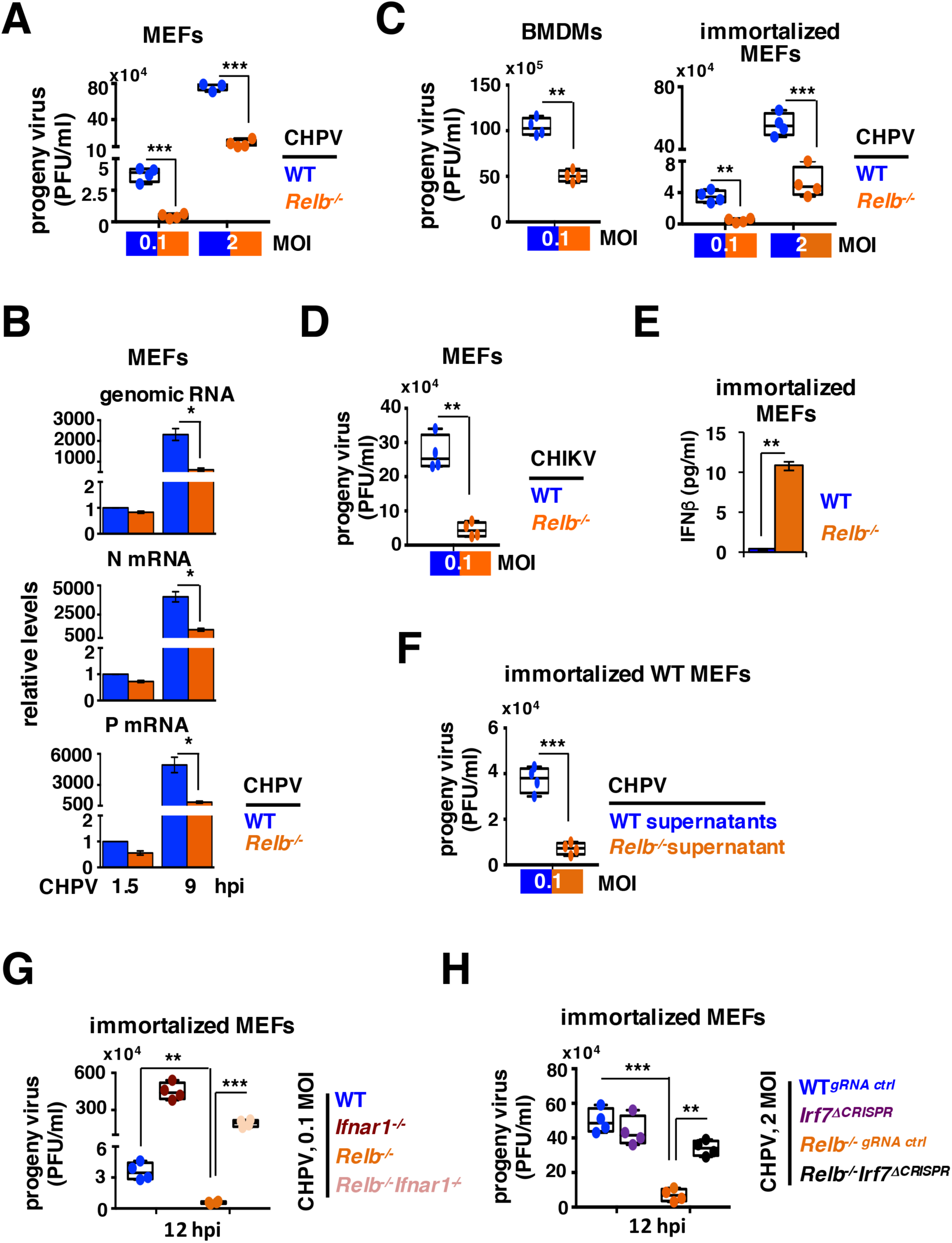
T1-IFNs and IRF7 cooperatively restrict viral multiplication in *Relb*^*-/-*^ cells. **A**. and **C**. Plaque assay revealing the titer of progeny virus particles produced by WT and *Relb*^*-/-*^ cells, including primary MEFs **(A)** or BMDMs **(C, left panel)** or immortalized MEFs **(C, right panel)**, upon infection with the indicated MOI of CHPV. **B**. qRT-PCR showing the relative abundance of viral genomic RNA, and mRNAs encoding viral Nucleocapsid protein and Phosphoprotein in WT and *Relb*^−/−^ MEFs at 9h post-infection with 2 MOI of CHPV. **D**. WT and *Relb*^*-/-*^ primary MEFs were infected with 0.1 MOI of CHIKV and the progeny virus titer in the culture supernatant was determined. **E**. ELISA quantifying the basal level of IFNβ present in the culture supernatants from immortalized WT and *Relb*^−/−^ MEFs. **F**. Immortalized WT MEFs were incubated with the culture supernatant obtained from immortalized WT or *Relb*^−/−^ MEFs for 8h before being subjected to 0.1 MOI of CHPV infections. The progeny virus yield was determined at 12h post-infection. **G**. and **H**. Plaque assay showing the tire of progeny virus particles produced by immortalized MEFs of the indicated genotypes upon infection with the indicated MOI of CHPV for 12h. The RT-qPCR and ELISA data represent means ± SEM of three independent experiments. The plaque assay data represent means ± SEM of four biological replicates. *, *P* ≤ 0.05; **, *P* ≤ 0.01; ns, not significant.

Our ELISA data further demonstrated that while almost undetectable in the WT supernatants, approximately 10 pg/ml of IFNβ was basally present in the culture supernatants derived from immortalized *Relb*^*-/-*^ MEFs (Figure 5E). Interestingly, incubating WT cells with the culture supernatant from immortalized *Relb*^*-/-*^ MEFs was sufficient to render those cells less permissive for CHPV multiplication (Figure 5F). Furthermore, compound genetic deficiency of IFNAR rescued CHPV growth in RelB-null cells (Figure 5G) while CRISPR-Cas9-mediated deletion of *Irf7* restored CHPV propagation even in the absence of RelB (Figure 5H). We conclude that RelB deficiency incites basal T1-IFN-IRF7 signaling instilling an altered cellular state restrictive to viral propagation.

### Vitamin D regulates antiviral type 1 IFN signaling *via* RelB and IRF7

We finally set out to address if RelB and IRF7 caused upregulation of antiviral T1-IFN signaling in vitamin D conditioned cells. Our mRNA analyses corroborated that vitamin D treatment significantly augmented the level of IRF7 mRNA in WT MEFs (Figure 6A), but did not impact the elevated basal expression of IFNβ as well as IRF7 mRNAs in *Relb*^*-/-*^ MEFs (Figure 6A and Figure 6B). Importantly, vitamin D conditioning was ineffective in further reducing the progeny virus titer in *Relb*^*-/-*^ MEFs infected with CHPV (Figure 6C). More so, abrogating IRF7 expressions in WT cells not only prevented heightened IFNβ mRNA synthesis upon vitamin D treatment (Figure 6D) but also obliterated the antiviral effect of vitamin D conditioning (Figure 6E). Our results suggest that RelB depletion upon vitamin D conditioning triggers the autoregulatory T1-IFN-IRF7 signaling circuitry, which reinforces cellular resistance to viral infections. In other words, RelB appears to instruct the antiviral innate immunity by modulating the homeostatic state of autoregulatory T1-IFN-IRF7 signaling (Figure 7).

**Figure 6:**
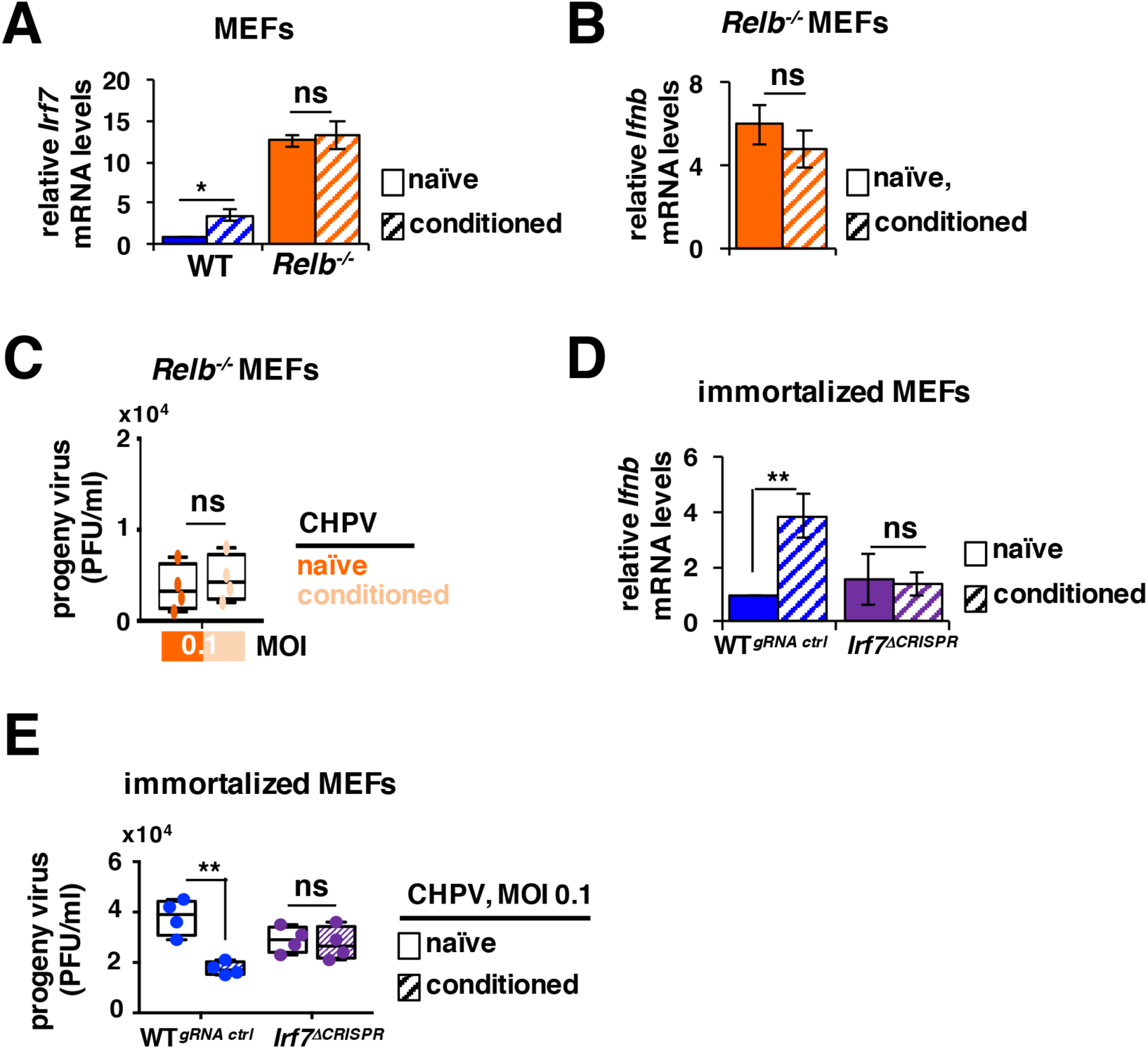
Vitamin D modulates the type 1 IFN-IRF7 pathway through RelB to exert antiviral effects. **A. B**. and **D**. Vitamin D naïve or vitamin D-conditioned WT or *Relb*^*-/-*^ primary MEFs were examined for the relative abundance of IRF7 mRNA **(A)**. Similarly, vitamin D-induced expressions of IFNβ mRNA were scored in *Relb*^*-/-*^ MEFs **(B)** and in immortalized *IRF7*^Δ*CRISPR*^ MEFs **(D)**. **C**. and **E**. Virus propagation was scored in vitamin D naïve and vitamin D-conditioned *Relb*^*-/-*^ MEFs **(C)** or immortalized *IRF7*^Δ*CRISPR*^ MEFs **(E)** subsequent to cell infection with 0.1 MOI of CHPV. The RT-qPCR data represent means ± SEM of three independent experiments. The plaque assay data represent means ± SEM of four biological replicates. *, *P* ≤ 0.05; **, *P* ≤ 0.01; ns, not significant.

**Figure 7:**
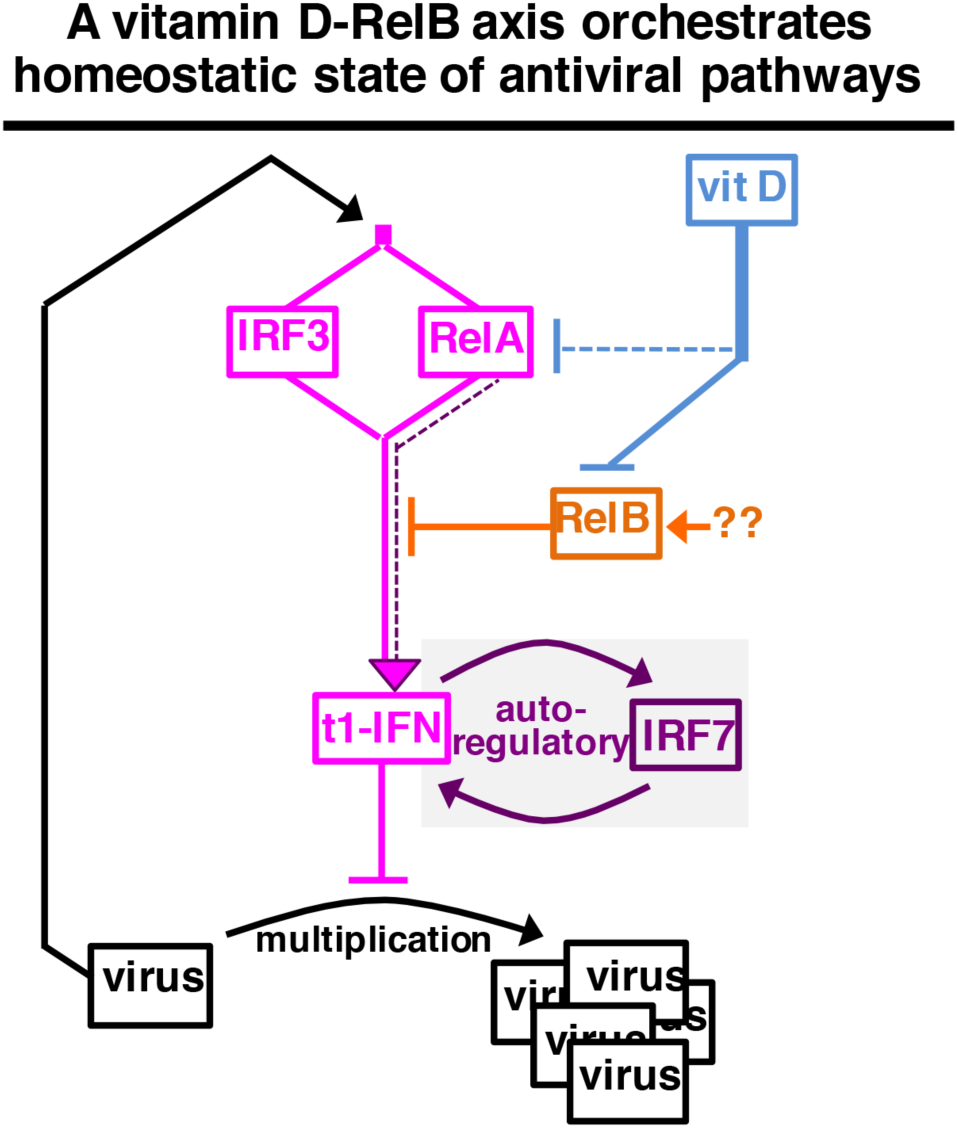
A schematic describing vitamin D-mediated modulation of the homeostatic state of antiviral signaling pathways.

## Discussion

Despite its well-articulated role in mitigating viral-inflicted pathologies, the molecular mechanism underlying antiviral vitamin D function remains obscure. Our CHPV infection studies unraveled a complex immune signaling circuitry restricting viral multiplications in vitamin D-exposed cells (Figure 7). The nuclear abundance of RelA and RelB largely determines the NF-κB transcriptional output. Based on the measurement of nuclear p52, Jin et al., (2014) earlier proposed that viral sensing activates the noncanonical RelB NF-κB pathway. We charted the NIK activity in our kinase assay and also scored the abundance of cellular p52 relative to p100 as a surrogate of noncanonical signaling. These direct measurements mostly ruled out the engagement of noncanonical signaling, at least in CHPV-infected MEFs. Of note, it was reported that a small fraction of RelB escapes p100-mediated inhibitions translocating to the nucleus independent of noncanonical signaling and that heightened RelB synthesis promotes such atypical RelB activation (Mineva et al., 2007; Roy et al., 2017). Our results indicated that the rate of RelB synthesis, more so than the noncanonical pathway, instructed the constitutive as well as virus-induced nuclear activity of RelB. Moreover, vitamin D conditioning severely impacted the abundance of RelB, both constitutive and virus-induced, in the nucleus by suppressing RelB transcription. Our genetic analyses strongly supported that RelB downregulation upon vitamin D conditioning provoked the autoregulatory T1-IFN-IRF7 circuitry, which strengthened antiviral cellular machinery. However, the observed vitamin D effects were moderate. We argue that such controlled modulation eluded toxicity associated with overtly active T1-IFN signaling while ensuring some protection from viral infections.

The *Relb* promoter possesses a κB site, which directs RelA driven transcription, as well as a vitamin D receptor element (VDRE), which instructs vitamin D sensitive gene expression (Bren et al., 2001; Dong et al., 2003). Indeed, it was shown that the canonical RelA/NF-κB pathway is important for both basal as well as cytokine-induced expressions of RelB mRNA (Basak et al., 2008; Chatterjee et al., 2019). In contrast, vitamin D treatment caused Histone deacetylase 3 dependent transcriptional repression of *Relb* in dendritic cells (Dong et al., 2005). These studies and our own led us to contend that the canonical RelA/NF-κB pathway promotes nuclear RelB by supporting RelB synthesis both in uninfected and CHPV-infected cells and that vitamin D acts through the VDRE present in the *Relb* promoter counteracting RelA-driven synthesis leading to depletion of cellular and nuclear RelB. Consistent to previous reports (Chen et al., 2013), vitamin D also inhibited canonical RelA/NF-κB signaling in our studies. We do not rule out that inhibition of RelA, in part, might have contributed to the downmodulation of RelB in vitamin D treated cells. However, we could conveniently establish that compared to RelA, vitamin D more drastically impacted the nuclear RelB activity with important functional consequences for T1-IFN expressions.

How does RelB control T1-IFN expressions? It was demonstrated that RelB binding displaces RelA from the *Ifnb* promoter leading to altered histone modifications and transcriptional repressions (Jin et al., 2014). In our experiments, RelB-deficiency intensified the constitutive synthesis of IFNβ as well as IFNα4 mRNAs. Albeit at a low level, RelA was shown to drive early IFNβ expressions in vesicular stomatitis virus-infected cells independent of IRF3 (Wang et al., 2010). Curiously, *Relb*^*-/-*^ cells displayed basal RelA, but not IRF3, activity in the nucleus. Our genetic studies indeed confirmed that canonical NF-κB subunits, including RelA, were required for the increased IFNβ production in RelB-null cells. Remarkably, autocrine IFNAR signaling and IRF7 were also necessary for maintaining the elevated IFNβ mRNA level in *Relb*^*-/-*^ cells. Except for plasmacytoid dendritic cells (pDCs), IRF7 is typically negligibly expressed in other cell types (Ning et al., 2011). Basally elevated IRF7 expression, in fact, enables pDCs to play a dominant T1-IFN-producing role in physiological settings. On the other hand, IFNβ synthesized early during viral infections engages autocrine IFNAR signaling, which leads to delayed accumulation of IRF7 in virtually all cell types (Sato et al., 1998). Newly synthesized IRF7, in turn, supports late T1-IFN synthesis reinforcing adaptive immune responses. Our studies captured the role of RelB in determining the homeostatic state of this autoregulatory T1-IFN-IRF7 signaling circuitry. In a mechanistic model (Figure 7), we reconcile that reduction of RelB, such as those achieved upon vitamin D treatment, primes RelA-driven synthesis of IFNβ, which signals through IFNAR inducing IRF7 production even in the absence of viral infections. IRF7 accumulated in RelB-depleted cells then sustains augmented expressions of not only IFNβ but also IFNα4 mRNAs. Importantly, our model readily explained why RelB deficiency rendered redundant, while IRF7 removal abrogated antiviral vitamin D actions.

Inadequate knowledge of the CHPV proteome and insufficiently characterized macromolecular interactions driving CHPV replication present serious challenges for the rational designing of antivirals (Chakraborty et al., 2021; Sharma et al., 2021). Our work mechanistically links vitamin D with anti-CHPV immunity at the cellular level. We argue that the proposed link may significantly impact the public health policy managing CHPV outbreaks in India and other countries if found true. Indeed, the physiological significance of this pathway needs to be firmly established in experimental animal models subjected to altered dietary vitamin D supplementation. Global *Relb*^*-/-*^ mice are less suitable for studying viral infections because of the lack of secondary lymph nodes and multiorgan inflammation, and the use of cell-type specific *Relb* knockout mice should be contemplated (Sun, 2017). Human population surveys charting vitamin D status in a geographic locale affected by the CHPV will also be informative. Finally, although we tested our mechanistic model in the context of cell infection by CHIKV, future studies ought to more rigorously determine if the vitamin D-RelB pathway plays a generic antiviral role. In sum, micronutrients are important immunomodulatory agents, but their potential in alleviating threats from microbial agents has not been fully explored (Richardson and Lovegrove, 2021). We suggest that an improved mechanistic understanding of how micronutrients modulate immune pathways may provide persuasive scientific arguments for micronutrient supplementation as prophylactics or adjunct therapy in epidemics, particularly those caused by viral pathogens.

## Materials and Methods

### Cells and viruses

Primary MEFs were generated from WT and gene-deficient C57BL/6 mice following the guidelines of the Institutional Animal Ethics Committee of NII (approval no. #380/15). Crossbreeding *Relb*^*-/-*^ and *Ifnar1*^*-/-*^ mice, composite *Relb*^*-/-*^*Ifnar1*^*-/-*^ MEFs were derived. As described earlier (Bais et al., 2019), these MEFs were used either as primary cells or subsequent to their immortalization by the NIH-3T3 protocol. BMDMs were generated using M-CSF following the published protocol (Manzanero, 2012). The CHPV strain 653514 was from the National Institute of Virology, Pune. The CHIKV strain 119067 has been described earlier (Bais et al., 2019). These viruses were propagated in VERO cells. Immortalized *Relb*^*-/-*^*Nfkb2*^*-/-*^ MEFs expressing RelB from a retroviral transgene under a promoter consisting of five tandem κB sites have been described earlier (Roy et al., 2017).

### Infection studies and cell treatments

WT or knockout cells were infected in serum-free media with CHPV or CHIKV for 1.5h at the indicated MOI. Subsequently, cells were washed and the culture was replenished with DMEM containing 10% serum. For vitamin D conditioning, cells were treated with 1α, 25-dihydroxyvitamin D3 (Sigma Aldrich) for 24h, and then infected with virus either subsequent to vitamin D withdrawal or in the continuing presence of vitamin D. At various times post-infection, the culture supernatant was collected and infected cells were harvested. As described (Bais et al., 2019), the abundance of virus particles in the supernatant was scored by plaque assay. Culture supernatants were also subjected to ELISA using the mouse IFN-β ELISA kit from PBL Assay Science, NJ.

### Biochemical studies

Whole-cell extracts and nuclear extracts, prepared from harvested cells, were subjected to immunoblot analyses and EMSA, respectively (Bais et al., 2019). NF-κB related antibodies have been described earlier (Mukherjee et al., 2017). Antibodies against p-IRF3 (#4947), IRF3 (#4302), NIK (#4994), TBP (#44059) and GAPDH (#2118) were purchased from Cell Signaling Technologies, USA. Anti-IRF7 antibody (AHP-1180) was from Bio-Rad laboratories. The kinase assay was performed following the previously published protocol (Mukherjee et al., 2021). The gel images were acquired using PhosphorImager (GE Amersham, UK) and quantified using ImageQuant 5.2.

### Gene expression studies

Total RNA was isolated from MEFs using RNeasy Kit (Qiagen), and RT-qPCR analyses were performed for determining the abundance of viral genomes and viral-encoded as well as host-derived mRNAs, as described earlier (Bais et al., 2019). Supplementary Table 1 provides a description of primers used in RT-qPCR studies. The microarray experiment was performed at Sandor Pvt. Ltd (Hyderabad, India) using Illumina MouseRef-8 v2.0 Expression BeadChip. We considered genes that have a detection p-value <0.05 in replicate samples and a signal to noise ratio, measured as the ratio of the average expression value to the standard deviation, ⩾2 across genetic background. Accordingly, we arrived onto a list of 8113 genes, which were then ranked in descending order based on the fold change difference in their basal expressions between *Relb*^*-/-*^ and WT cells. The relative enrichment of T1-IFN pathway genes in this ranked list was determined by GSEA using fgsea package in the R platform (Subramanian et al., 2005); T1-IFN pathway geneset was from the Reactome Pathway database with an ID - R-HSA-909733. The MIAME version of our microarray data is available at NCBI-GEO (accession no. GSE176446). For investigating the vitamin D-activated gene program, we examined a previously published transcriptomics dataset obtained using THP1 human-derived monocytic cell line (NCBI-GEO, accession no. GSE57028) (Salamon et al., 2014). Correcting for the detection p-value and signal to noise ratio, we prepared a list of 27164 genes that was subjected to GSEA.

### Generation of ΔCRISPR cell lines

Lentiviral particles encoding guide RNA and Cas9 were produced in HEK 293T cells using mouse *Irf7* CRISPR lentiviral transfer construct targeting exon 7 (target ID #MM0000202587) of *Irf7* (Sigma Aldrich). Lenti CRISPR universal non-target plasmid was used as a control (Sigma Aldrich). WT or *Relb*^*-/-*^ immortalized MEFs were then transduced with lentivirus and transduced cells were FACS sorted and further selected using puromycin. The efficacy of CRISPR/Cas9 genome editing in the resultant *Irf7*^Δ*CRISPR*^ and *Relb*_*-/-*_*Irf7* ^Δ*CRISPR*^ cells were evaluated in immunoblot analyses. Supplementary Table 2 provides a description of gRNA used in this study.

### Statistical analysis

Error bars are presented as standard errors of the means (SEM) of three or more experimental replicates. The statistical significance was calculated using two-tailed Student’s t-test. * signifies P ≤ 0.05, ** and *** denotes P ≤ 0.01 and P ≤0.001, respectively.

## Supplemental Information

Supplemental Information includes one figure and two tables associated with the main text.

## Acknowledgement

We sincerely thank Dr. P. Nagarajan, SAF for the help with animal husbandry and V. Kumar, SIL for technical assistance. We also thank the members of SIL for help and suggestions. Virus research in the PI’s laboratory is funded by the S. N. Ramachandran National Bioscience Award to S.B by Department of Biotechnology, Govt. of India and support from NII-core. Y.R. thanks the Indian Council of Medical Research, India for research fellowship.

## Author Contribution

YR carried out the wet laboratory experiments with the help from NK, CB, and SSB. GAA guided experiments involving vitamin D. Transcriptomics data analyses were performed by MKS with the help from CC and GAA. CHIKV experiments were guided by GM. S.B. conceived and supervised the overall research and wrote the manuscript with YR. The authors declare that they have no conflicts of interest with the contents of this article.

