## Supplemental Data for "A vitamin D-RelB/NF-κB pathway limits Chandipura virus multiplication by rewiring the homeostatic state of autoregulatory type 1 interferon-IRF7 signaling"

**# Figure S1;**
**# Supplementary Table S1-S2;**

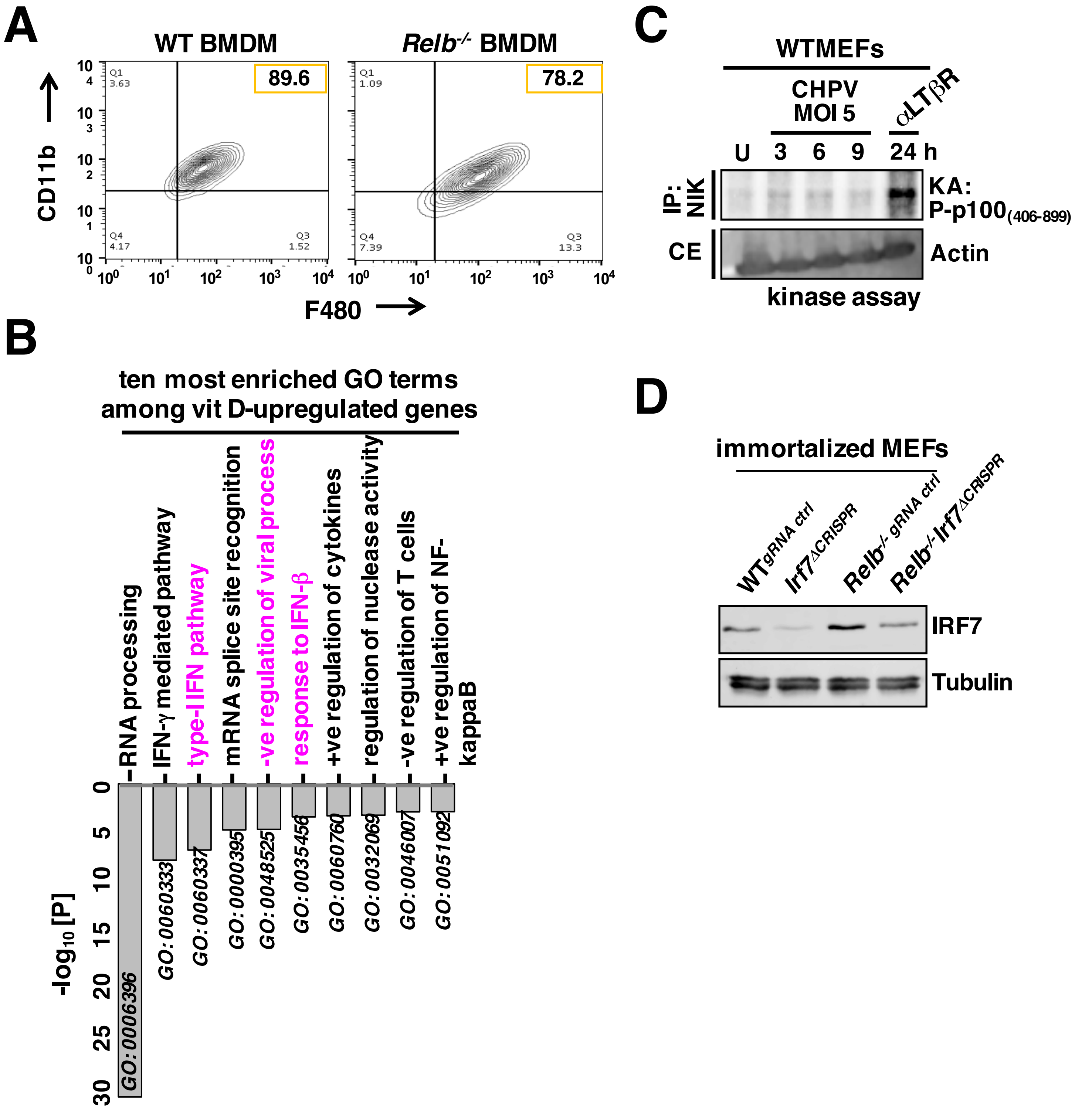


**Figure S1: Investigating the vitamin D-RelB signaling axis in cultured cells. A.** Flow cytometry analyses showing the purity of macrophage population differentiated ex vivo from bone marrow cells. **B.** Enrichment analyses identifying the top ten most enriched GO for biological processes terms among vitamin D-upregulated genes in THP1 cells. Briefly, published dataset with NCBI-GEO accession no. GSE57028 was used. Considering genes that were induced at least 1.5 fold upon 24h of vitamin D treatment in replicate samples with a signal-to-noise ratio ⩾2, we arrived onto a list of 853 genes. We subjected the gene list to GO analyses. The entire Illumina MouseRef-8 v2.0 gene-array was used as the background. Corresponding enrichment scores have been plotted and GO IDs have been indicated. GO terms linked to the antiviral type-1 interferon pathway has been indicated in magenta. **C.** Kinase assay revealing the activity of NIK in the indicated cell extracts. Cells treated with agonistic αLTβR antibody has been used as a positive control. **D.** Immunoblot analyses assessing the efficiency of gRNA mediated depletion of IRF7 in immortalized WT or *Relb*$^{-/-}$ MEFs.

**Supplementary Table S1: primers for RT-qPCR**

| Gene Name | Forward 5'→3' | Reverse 5'→3' |
|---|---|---|
| **CHPV Genome** | CGAGTGAACTCAGTTGCAGAG | GAATCGAGAGTGTCCTGAAGC |
| **CHPV N** | GATTTGTTGCGGATGATGAC | CCAGAAATGGAAACTGGGAT |
| **CHPV P** | CTCTCCGTCTGATCCACCTT | TCAATCCAGCAATGACCAGT |
| **mIFNβ** | CCGGACTTCAAGATCCCTATGGA | TGGCAAAGGCAGTGTAACTCTTC |
| **mIFNa4** | CCTGTGTGATGCAGGAACC | TCACCTCCCAFFCAGTGA |
| **ISG56** | TGGCCGTTTCCTACAGTT | TCCTCCAAGCAAAGGACTTC |
| **ISG15** | AGCTCCATGTCGGTGTCAG | GAAGGTCAGCCAGAACAGGT |
| **OAS1** | AGGGGCATTTGCTGCTCTGC | GGGCACCTGCTGTGGTTTATTG |
| **IRF7** | CTGGAGCCATGGGTATGCA | AAGCACAAGCCGAGACTGCT |
| **IL-1b** | AACCTGCTGGTGTGTGACGTTC | CAGCACGAGGCTTTTTTGTTGT |
| **RelA** | GCAGTATCCATAGCTTCCAG | TGCTCCTCTATAGGAACGTG |
| **RelB** | GAATGTCGTCAGGATCTGC | TGGTGGACTTCTTGTCGTAG |
| **mActin** | CCAACCGTGAAAAGATGAC | GTACGACCAGAGGCATACAG |

**Supplementary Table S2: descriptoin of gRNAs used in this study**

| Target ID | Target Sequence 5'→3' | PAM | Target gene symbol |
|---|---|---|---|
| **MM0000202587** | ATGTGACCATCATGTACAA | GGG | Irf7 |
